# A ground surface rolling method for detecting environmental DNA of terrestrial animals

**DOI:** 10.1101/2024.09.19.613995

**Authors:** Manami Kakita, Yoshikazu Furuta, Hidenori Tanaka

## Abstract

Biological information obtained from environmental DNA has been helpful for conservation of ecosystems and biodiversity. Methods for environmental DNA of terrestrial habitats are limited such as sampling of water near terrestrial habitat and of shoveled soil, whereas those of aquatic habitats are well-established. We developed a method using a sampler for terrestrial habitat utilizing a combination of a simple rotating body and a single-use non-woven fabric collector as a sampler, enabling sample collection from wide area of various surfaces without power supply. The sampler was applied to collect 90 ground surface and 29 water samples in a secondary nature environment, resulted in the detection of 53 species of birds and mammals in total. The result covered 92% of the species detected by cameras installed in front of the sampling areas, validating the sensitivity of the method. Time course analysis also revealed that the timing of the detection of birds matched that of the birds’ actual arrival. These results suggest that our labor-saving method could provide accurate and time-resolved data for biodiversity monitoring of terrestrial areas. Combining with established aquatic environmental analysis, the sampler would enable systematic monitoring of both aquatic and terrestrial habitats, enhancing our understanding of the whole ecosystem.

## Introduction

For ensuring humans live in harmony with nature, it is essential to monitor biodiversity by conducting and assessing conservation activities to understand the impact of anthropogenic activities on biodiversity. It is also necessary to understand the environmental conditions from detailed biodiversity information to consider policies, activities, and decisions whether human should intervene in the environment for biodiversity conservation.

A widely used biomonitoring method is morphological surveys such as visual observation, capture surveys, and camera traps, but these methods are labor intensive and require long survey times in the field. Recently, environmental DNA analysis has become commonly used as a novel monitoring method^1–4^. Environmental DNA is biogenic DNA present in environmental media such as water, soil, and air. It is released from living organisms through feces, skin fragments, body fluids, and dead bodies and accumulates in the environment^5^. Aquatic life monitoring using environmental DNA sampled from waterbodies has standardized protocols and are commonly used for habitat surveys mainly in rivers and oceans^6–8^. As it can be conducted only with field sampling and does not necessarily require expertise as in trapping surveys, visual observations, or species identification, the method is widely used by various sectors such as academic, governmental, and business.

In contrast, environmental DNA analysis techniques for terrestrial habitats including forests and urban areas are less developed compared to that for aquatic habitats^9–11^. In recent years, there have been increasing reports on the use of environmental DNA analysis to detect terrestrial animals such as birds and mammals, mainly utilizing one or combinations of the three types of source materials: water, soil, and air. Water can be utilized to detect some terrestrial organisms, but the detection is limited to species which have direct contact with the water^12–15^. Soil samples shoveled from multiple locations can collect the terrestrial organisms derived environmental DNA more directly compared to water sampling, but large-scale sampling is challenging in terms of the labor effort required^16–19^. Also, as it is reported that DNA remains in the soil for longer periods compared to water^20–24^, acquired data would be a mixture of new and old contacts with organisms and care should be taken for data interpretation. Air sampling was also used to collect and detect terrestrial environmental DNA as it can collect diffused environmental DNA from wide area, but air samplers usually require a power source for air suction to collect samples, which could be a challenge for easy and convenient sampling^25–27^. Additionally, noise emission by air samplers would disturb wildlife. Furthermore, detection of animals of small populations might be difficult as environmental DNA sampled by air samplers usually results in quite low DNA concentration. Considering drawbacks of the methods and materials above, experimental protocols with a sampler that is power source-free system, easy to construct and handle, and applicable for sampling of wide area would be ideal for sampling of terrestrial animals derived environmental DNA.

Here, we developed a new protocol for sampling of terrestrial animals derived environmental DNA utilizing a rotating body and a single-use non-woven fabric collector. The sampler, hereafter Koro-rin sampler, is easy to construct and can be used without a power source. It can also easily and rapidly collect environmental DNA on various kinds of ground surface, not limited to soil but also applicable on rocky surfaces, hard ground surfaces, tree roots, grasses, and other ground surfaces that are difficult to shovel. Using the self-crafted Koro-rin sampler, we conducted weekly sampling of a terrestrial area of a secondary nature environment for 14 months, followed by metabarcoding analyses for birds and mammals. The main objectives of this study were (1) to demonstrate the ability of the sampler to detect terrestrial animals, (2) to validate the sensitivity of the sampler for species identification by comparing the results with camera trap data, and (3) to verify whether the sampler is applicable for time-course analysis of environmental DNA on the ground surface. The results provided insights on the applicability of surface sampling for biodiversity monitoring of terrestrial habitats.

## Results

### Construction and usage of Koro-rin sampler

For easy and efficient sampling of various surfaces of terrestrial habitats, we developed a sampler utilizing a rotating body and a single-use non-woven fabric collector, named Koro-rin sampler. The use of a single-use non-woven fabric for the surface, rather than the sticky sheet commonly used as cleaning apparatus, allows to trap sediments on the ground surface more efficiently. We also added the cushioning of the rotating body so that it can be efficiently used on non-flat surface. With holding and pushing a long edge, a person can use the sampler in a standing position to collect a wide area of the sampling site of interest.

For the validation of the sampler for the detection of terrestrial species, ground surface and water samples were collected using the sampler from November 2021 through December 2022 at the Forest of Toyota in Toyota City, Aichi Prefecture, Japan (Fig. 1a). The samples were weekly collected from four sites: three soil-exposed ground surface sites (Sites 1-3) by using the sampler and one water point with flow (Site 4) by sampling water body. A total of 90 surface samples and 29 water samples were collected during the sampling period and species were detected using metabarcoding analysis using MiBird and MiMammal primers^12,28^.

**Fig. 1:**
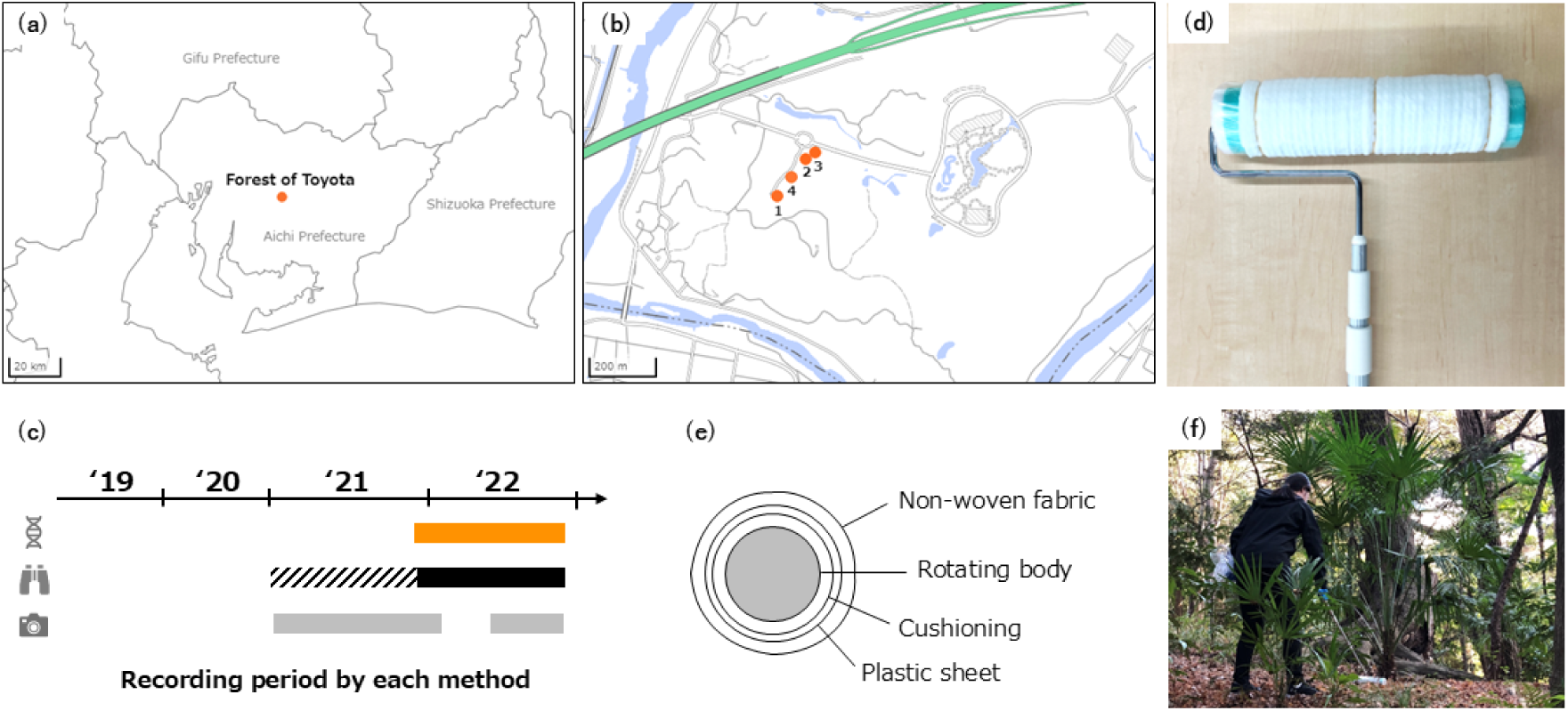
Location and duration of environmental DNA sampling, and images during sampling. **a** Location of Forest of Toyota (Iwakura-cho, Toyota City, Aichi Prefecture, Japan) where sampling was conducted. **b** Detailed locations of the four sampling sites. Sites 1-3 were sampled using an Koro-rin sampler to collect about 10 m^2^ of ground surface, and site 4 was sampled using a syringe to collect250 ml of environmental water. These maps were generated using the GSI map Vector (https://maps.gsi.go.jp/vector/ Accessed June 19, 2023). **c** The periods of record for DNA, camera, and visual surveys are shown below; DNA was conducted from November 11, 2021, to December 23, 2022; trail cameras were conducted from January 26, 2021, to December 23, 2022 (excluding January 27-April 6, 2022), and visual surveys were conducted from 2021-2022. See the main text for details. **d** Image of the Koro-rin sampler. **e** Sampler components and cross-sectional view. **f** Image of ground surface sampling using the Koro-rin sampler.

As some species apparently exist in the Forest of Toyota but lacked in the reference sequence database, we determined the sequence of the region of MiBird and MiMammal of several animals using its feces or feather specimen. Obtained sequences were manually added to the database and used for the species detection (See Materials and Methods for more detail).

### Samples collected using Koro-rin sampler resulted in the detection of 53 wild species

First, we tested how many species can be detected using the Koro-rin sampler. After aggregating the obtained data at the genus level and excluding genus obviously derived from domesticated animals, 53 genera (30 birds and 23 mammals) were detected more than once among the samples of all the timepoints of the ground surface (Table 1). Comparing genus detected from ground surfaces and water, all the genera detected in the water samples were found at one or more of the three ground surface sites (Supplementary Table S1). Additionally, 18 of the 30 wild bird genera (60%) and 15 of the 23 wild mammal genera (65%) were detected only from the ground surface samples. Addition of genus detected from the ground surfaces to those from water resulted in two-fold increase of the number of detected genera, suggesting wider coverage of the terrestrial biodiversity by the ground surface samples than water samples. Regarding species detected from both water and ground surfaces, small bird groups that congregated at water sites, including *Parus* and *Sittiparus*, were detected more frequently in water than on ground surfaces.

**Table 1:**
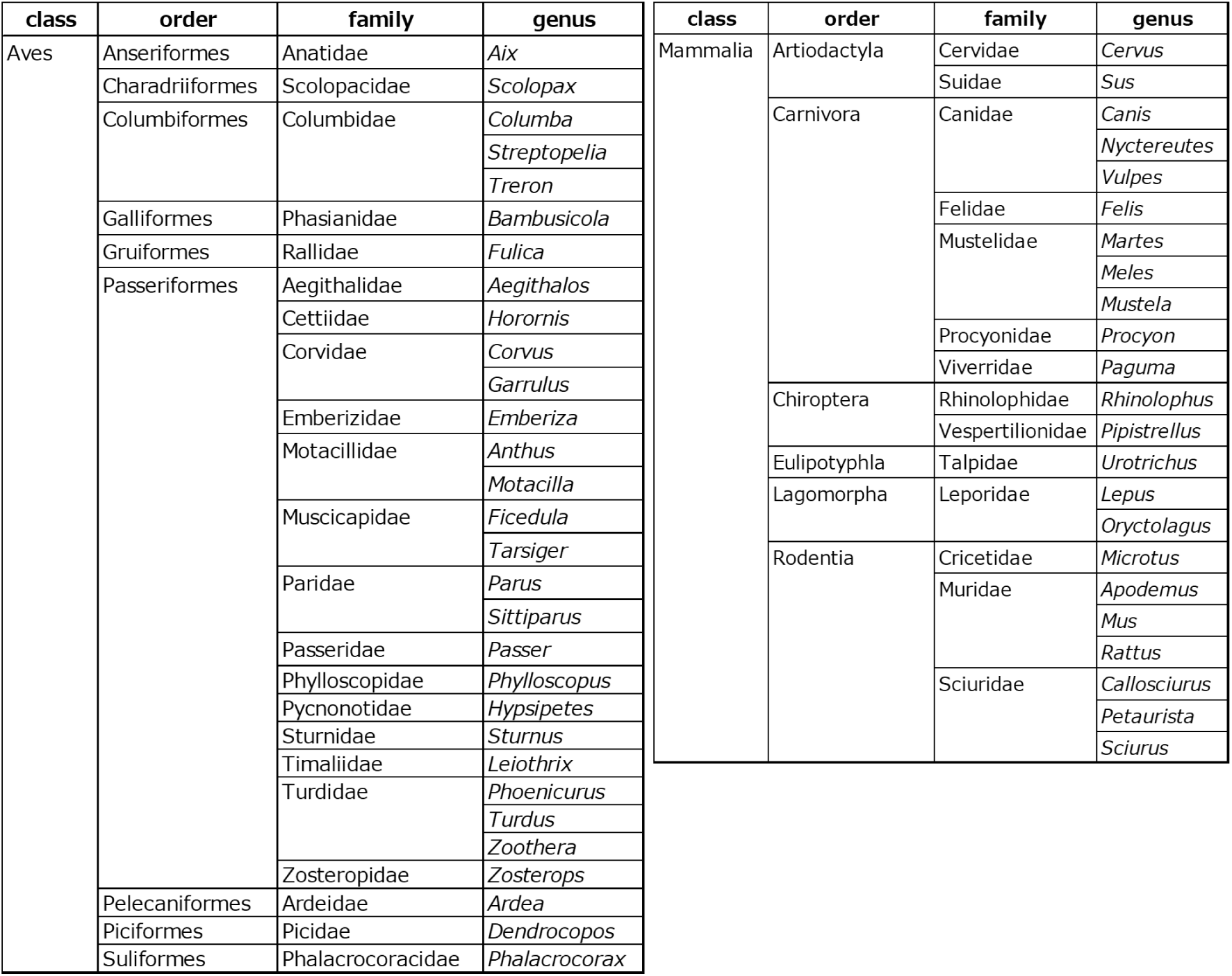
List of birds and mammals detected in the metabarcoding analysis.

Results of the metabarcoding was also compared between the sampling sites and detected, 21, 20, 22, and 12 bird species and 20, 19, 17, and 8 mammal species at the ground surface sampling sites 1, 2, and 3 and the water site, respectively (Fig. 2). The variation among sites was particularly high for birds, with 12/30 genera detected at all sites or at three ground surface sites and 18/30 genera detected at one or two sites. This would be because once environmental DNA is attached and deposited on the ground surface, it does not diffuse much and tends to remain at the same place.

**Fig. 2:**
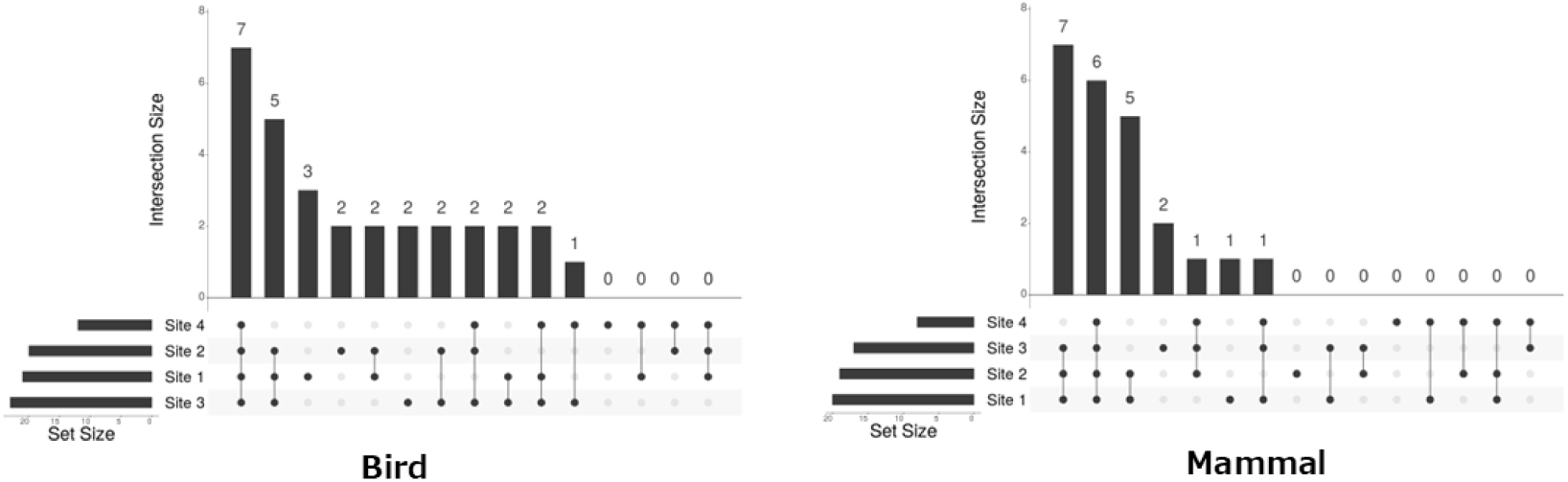
Variation in DNA detection species of each bird and mammal at the four sites. Detected species shared among each site are shown in Upset-Plot. Set Size in the lower left corner indicates the total number of species detected at each site, and Intersection Size indicates the number of species detected shared by each site combination.

### Degree of overlap between DNA and species sighted throughout the environment

To validate whether the genera detected by the sampler matches the genera were present, the results of the metabarcoding was compared with the data of visual surveys throughout the Forest of Toyota during the sampling period (Table 2). The visual surveys detected 65 genera (47 birds and 18 mammals) throughout the Forest of Toyota. Although only three small sites (Approximately 30 m^2^ in total) were applied for the ground surface sampling, the result of metabarcoding covered 43/65 (66%) of the genera detected in the visual survey throughout the whole area of the Forest of Toyota (Approximately 450000 m^2^). The 22 genera which were not detected in the metabarcoding analysis would be the genera which observed only at the sites distant from the sampling area and/or the genera with low abundance. In contrast, among the genera detected by the metabarcoding, 10/53 (19%) genera were only detected in the metabarcoding analysis. The 10 genera included 6 small animals which are usually difficult to observe visually, demonstrating the better sensitivity of the metabarcoding analysis for small animals.

**Table 2:**
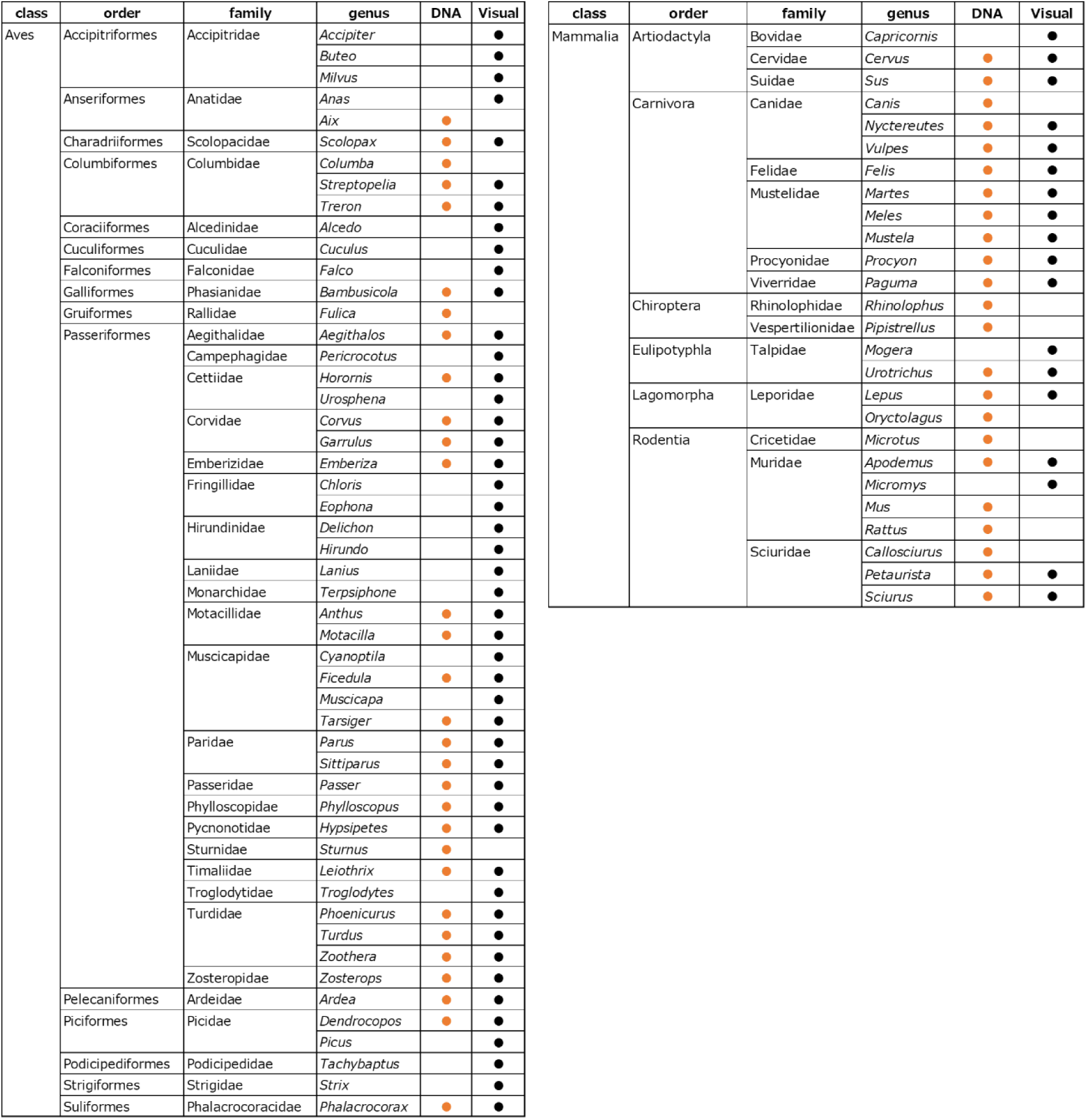
Overlap between birds and mammals visually observed during the sampling period and DNA detections.

### The Koro-rin sampler method covered 92% of the camera-detected genera

For the further validation of the sampler with visual data, the metabarcoding results were also compared with the data of trail cameras which were set just in front of the sampling sites (Fig. 3). The trail cameras captured animals passed the sites by motion detection between January 26, 2021, and December 23, 2022, which included the period before the sampling started. The captured data, accumulated as 3,092 images, were inspected by wildlife experts manually to identify bird and mammal genera. Twenty-six genera were captured by the trail cameras set up at all the ground surface sampling sites during this period, of which 24/26 (92%) overlapped with the genera detected by metabarcoding. The species that were only detected by the cameras but not by metabarcoding were *Strix* and *Capricornis* spp. *Strix* was recorded only once on October 18, 2021, which was before the sampling started (November 2021). *Capricornis* was recorded on April 12, 2021, April 24, 2022, and October 14, 2022, of which two recordings overlapped with the DNA sampling period. In contrast, a relatively low proportion of the metabarcoding-detected genera (24/53, 45%) were detected by the cameras. The 29 species that were detected only by the metabarcoding included many small mammals, such as bats and rodents, as well as birds that did not often land on the ground, which could be difficult to capture by the cameras targeting the ground. These results demonstrated the high performance of our method in the detection of terrestrial animals.

**Fig. 3:**
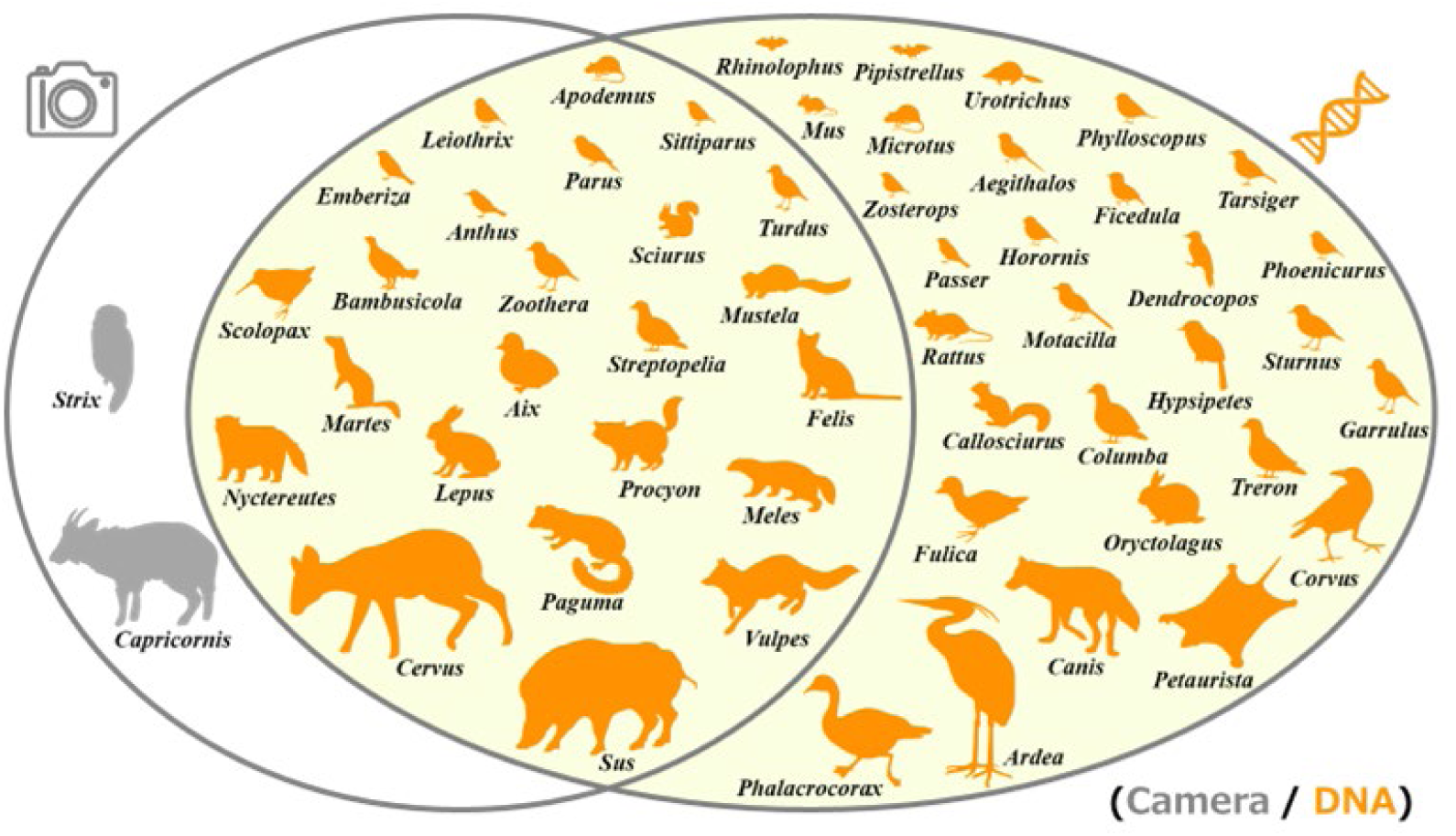
Degree of overlap between species detected by DNA and recorded by trail cameras. The results of bird and mammal species identification from 3092 images taken by trail cameras and the degree of overlap between the results of detection by DNA are shown. All results were aggregated at the genus level. Species that were only detected by camera were painted in gray and those detected by DNA were painted in orange.

### Time-resolved data acquisition

To determine whether the metabarcoding can detect the seasonal arrival and leave of birds, we investigated the perturbation of the detection of bird genera throughout the time course and compared with the timing of the observation of the genera by the visual survey throughout the Forest of Toyota, focusing on two resident bird species, a summer bird species, and a winter bird species as representatives (Fig. 4). The brown-eared bulbul (*Hypsipetes*) and warbling white-eye (*Zosterops*), the two resident bird species frequently observed throughout the year, were also detected throughout the year by the metabarcoding. On the other hand, the narcissus flycatcher (*Ficedula*), visually observed in the warm season, and white thrush (*Zoothera*), visually observed in the cold season, were also detected by the metabarcoding at the timepoints close to the visually observed timepoints. The results suggest that the analysis of the ground surface samples and the metabarcoding can detect the seasonal variation of the existence of birds. The detection of the genera was lost as the bird left probably because of the quick turnover of the bird derived DNA on the soil surface.

**Fig. 4:**
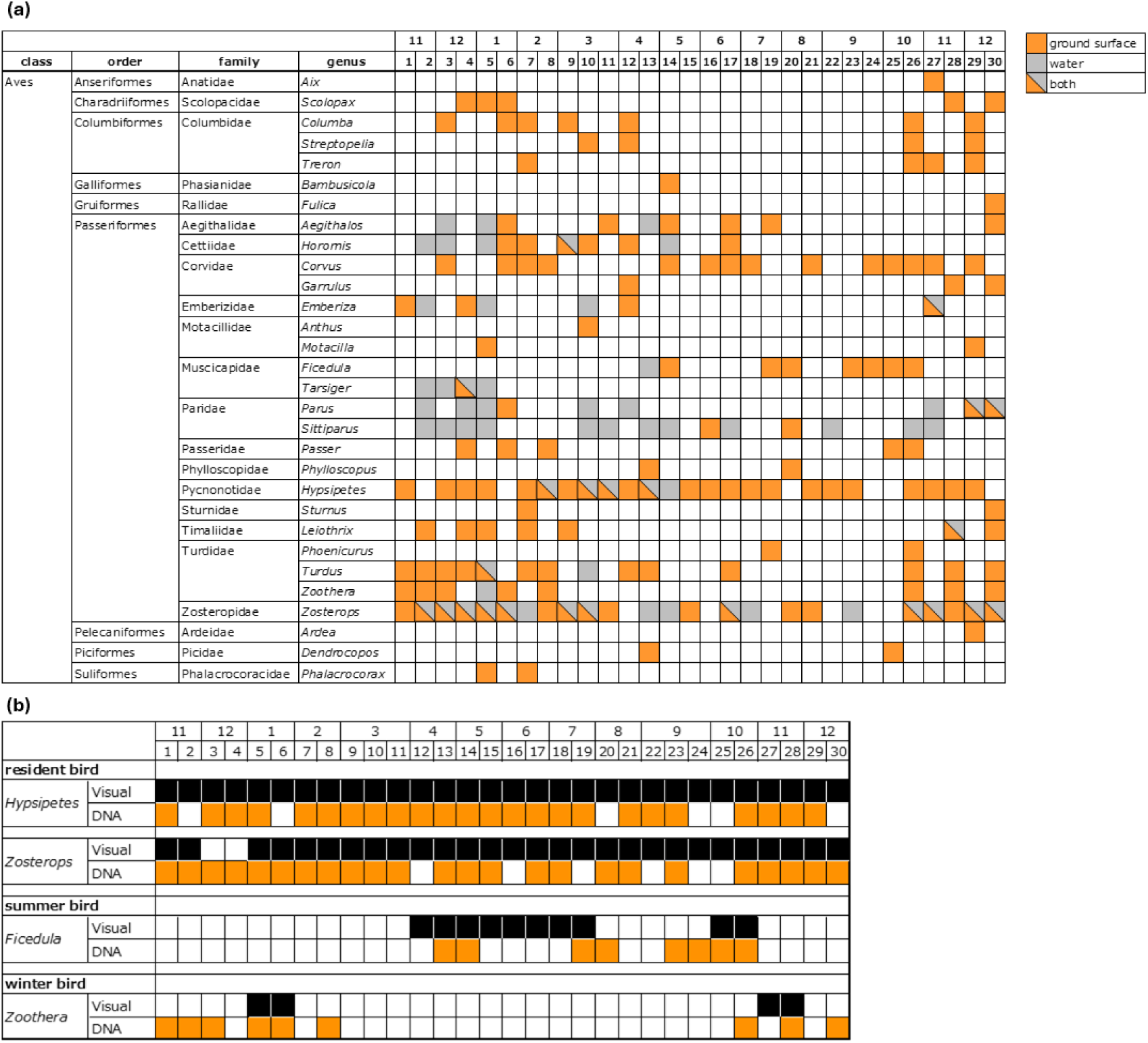
When each bird was detected by DNA. **a** A time-series plot of the bird species detected by DNA and the time points in which they were detected is shown. Numbers 1-12 indicate months and numbers 1-30 indicate sampling times. Detections from the ground surface only are shown in orange, those from environmental water only in gray, and those from both are separated by a shaded line and painted in both colors. **b** We focused on four representative resident, summer, and winter bird species of all birds detected by DNA, and compared the period when they were visually observed with the period when they were detected by DNA. The period of visual observation was painted in black, and the period of DNA detection was painted in orange.

We also compared the time course of the results of metabarcoding and the trail camera data, which included more observation timepoints than the visual survey and enabled the comparison on a per-sampling basis. For the 1st to 6th sampling time point, the metabarcoding results of a sampling time point were compared with the camera data during the period between the time point of interest and the previous sampling timepoint, which was between 13 to 15 days. For the 1st sampling, we used the camera data from the previous 14 days.

From the 1st to 6th sampling, the proportions of DNA detected relative to the camera data were 71%, 100%, 100%, 71%, 80%, and 38%, respectively (Fig. 5a). Comparison of the genera detected at each sampling timepoints suggested that the change of the genera captured by the camera was also covered by the metabarcoding (Fig. 5b). For example, Woodcock (*Scolopax*), which was detected exclusively in the 4th sampling timepoint by the camera, was also detected exclusively by the metabarcoding. These results totally indicated that the metabarcoding results can sensitively detect the change of the environmental DNA at the ground surface.

## Discussion

Using the Koro-rin sampler, the ground surface was sampled for environmental DNA to demonstrate the usefulness of the proposed method in biodiversity surveys. We found 53 wild bird genera and mammal genera in total by the metabarcoding analysis. The detected genera well covered those detected by visual survey and by trail cameras. The method also well captured the time course changes of the existence of animals, suggesting the usefulness of the method for environmental DNA monitoring in terrestrial environments.

Development of a method for sampling and analysis of environmental DNA in terrestrial environments is lagged from that for aquatic systems. One reason is the uneven distribution of environmental DNA in terrestrial environments^19,22,23^. In the case of aquatic systems, environmental DNA released from various sources such as feces, body fluids, and skin fragments diffuses in the water or transported by water flow. Therefore, biological information derived from a wide area can be possibly sampled in a single sampling^29–31^. However, in terrestrial environments, distribution of environmental DNA on the ground hardly diffuses nor distributes uniformly^22^. Therefore, compared to the sampling of aquatic areas, experimental conditions such as sampling sites and sampling methods should be carefully designed for the sampling of terrestrial environment to maximize the number of animal species detection.

Several approaches have been used to detect DNA from terrestrial animals in the environment; however, most of these approaches have been based on water or sediment sampling of waterbodies. In natural environments, the degree to which animal information can be determined from water is affected by the location of waterbodies, the amount and flow of water, and the animal behavioral routes^14,32^. Waterbodies in terrestrial environments often have low transparency and high turbidity^33^, which affect the amount of sample that can be filtered. Moreover, environmental DNA sampled from waterbodies might not fully reflect the whole biodiversity of the terrestrial animals when the animals congregate at waterbodies out of the survey area and/or when no information is available about the combination of animals and waterbodies they congregate. Our motivation was to maximize the detection of terrestrial species to obtain better snapshot of the biodiversity, thus we set out to construct a method for sampling the ground surface, which would complement or wholly include the species detected by water samples. Indeed, our analysis showed that the metabarcoding result of the ground surface samples wholly included the genera detected by water samples, demonstrating the necessity of the ground derived samples for the better coverage of the environment.

As our method developed in this work mainly collects soil samples, the results should be discussed with the previous works that sampled soil for the detection of terrestrial animals. An example of the comparison of genera detected from environment DNA sampled from soil and those from camera trapping was previously shown in^16^. Most of the mammals identified by camera were also detected by DNA. In addition, several small mammals that were not recorded by camera were detected by the metabarcoding analysis. These results are consistent with our results and with other studies that have mentioned the difficulty of capturing small mammals with cameras^34,35^, suggesting that environmental DNA surveys are effective for detection of small mammals.

Although our results showed similar sensitivity for small mammals with previous works that analyzed soil, the method for the sampling was totally different. The experiment conducted in the previous study^16^ used a protocol for collecting 2 cm scoops of surface soil at every 2 m, with 10 g used for analysis in 1 L, and reported that camera recordings taken 30–150 d prior to sampling were well reflected in the detected DNA. Other studies have indicated that the deeper the soil layer, the longer the environmental DNA in the soil is preserved^22^. If the purpose is to collect relatively old animal tracks, sampling should be expanded in the sub-surfaces and at the surface. In contrast, the sampler we developed samples only the ground surface, which greatly simplified the sampling process and also the results are supposed to detect the species that existed in the environment in a narrower time range by avoiding the collection of remaining DNA derived from old contact.

The sampler used in this study has a simple structure comprising a rotating body and single-use non-woven fabric collector. As the sampler has a simple configuration that can be easily crafted, application of this new sampling method can be adopted with little effort. It can collect samples through rolling, which is quicker and easier compared to other soil shoveling based methods. Because samples can be collected without shoveling the ground, the sampler can be used on rocky surfaces, artificial ground surfaces such as asphalt and concrete, tree roots, lawns, and other ground surfaces that are otherwise difficult to shovel. Another important feature of the sampler is that it does not require a power source or any additional equipment so that sampling is neither limited by power availability nor by the amount of battery power, which is different from the methods using air samplers. Furthermore, during the operation, the sampler creates extremely low noise and vibrations; thus, it does not disturb wildlife in natural habitats. It can also be mass-produced for large scale sampling because it is composed only of materials that can be easily procured at a low cost.

Although our method including the sampler was shown to detect eDNA derived from terrestrial animals, sensitivity can be further improved by resolving remaining issues specific to terrestrial samples. The sampler could capture larger amount of eDNA by considering the length of the rotating body and by considering the material used for the non-woven fabric collector. As soil includes large number of cells of microorganisms, a method that reduces microorganisms derived DNA during the DNA extraction process would improve the yield of animal derived DNA by avoiding non-specific amplification^6,36,37^. As rainfall can affect the results by physical washout and degradation of environmental DNA^38,39^, sampling day should be carefully chosen according to the weather if possible. Scarcity of the DNA sequence of terrestrial animals compared to fish is another problem for detection of wide variety of terrestrial animal species by metabarcoding^8,40^. As such, improvement from the various aspects of the methods should be accomplished for better detection of terrestrial animals by metabarcoding.

In conclusion, we proposed a simple environmental DNA collection tool, the Koro-rin sampler, for the detection of terrestrial animals and demonstrated its usefulness by monitoring wildlife and sampling their DNA in a secondary natural environment. We found that our method covered 66% of the animal genera visually observed throughout the environment and 92% of the genera detected by the trail cameras at the sampling sites. The data also showed that it can detect small mammals which are usually difficult to visually observe and that it can detect arrival and leave of specific birds in a timely manner. These results suggested that sampling of environmental DNA attached to and deposited on the ground surface is effective for detecting terrestrial wildlife in the environment. This method can be used by itself or with combination of other methods for biodiversity monitoring to capture the biodiversity of the whole ecosystem.

## Methods

### Wildlife monitoring using trail cameras and visual surveys

All field surveys were conducted in the Forest of Toyota (Toyota City, Aichi Prefecture, Japan) owned by the Toyota Motor Corporation (https://www.toyota.co.jp/jpn/sustainability/social_contribution/feature/forest/forest_of_toyota/). The forest is a 45 ha secondary forest (Satoyama) mainly consisting of *Quercus serrata* (jolcham oak) and *Quercus variabilis* (Chinese cork oak), which is well maintained and utilized for environmental education and morphological surveys. In visual surveys, species were sighted throughout the Forest of Toyota during the sampling period. Mammals were surveyed by year, so data for 2021-2022 (shaded and black areas) were used; birds were surveyed by month, so data from November 2021 to December 2022(black areas) were used (Fig. 1c). In trail camera surveys, among several trail cameras installed in the Forest of Toyota for wildlife monitoring, we collected captured images from trail cameras at three sampling locations to identify the species passed by. The 3,092 images captured between January 26, 2021, and December 23, 2022, including the period before sampling were manually inspected by a wildlife expert for identification of birds and mammals and compared with the environmental DNA analysis results (Fig. 1c). Of note, cameras were malfunction due to lightning strikes during the period from January 27 to April 6, 2022.

### Koro-rin sampler

A Koro-rin sampler for collection of ground surface sediments containing environmental DNA was constructed by wrapping a plastic bag (250 mm× 350 mm) around the roller part of Korokoro® Flooring Cleaner Midori (Nitoms, Japan), followed by wrapping a non-woven fabric sheet (Quickle Wiper three-dimensional adsorption sheet, Kao, Japan), secured both termini using rubber bands. The sampler was sealed and stored in a plastic bag until further use. To use the sampler, the roller was lightly pressed against the ground surface and rolled onto a sheet to collect ground sediments. To prevent DNA contamination from an operator person, nitrile gloves were worn during the fabrication and usage of the sampler with changing gloves for each operation. For every sampling day, eDNA was isolated from sheets which were not used for the sampling and used as negative controls to validate the absence of DNA contamination from an operator and the absence of background detection of birds and mammals.

### DNA sampling

Three sampling sites for terrestrial samples were chosen in the Forest of Toyota so that the location was captured by a trail camera. Sampling was conducted every 13–15 d for 30 times in total from November 11, 2021 to December 23, 2022 (Fig. 1c). The sampling area per sampling site was approximately 10 m^2^. After sampling, the non-woven sheets were separated from the Koro-rin sampler and immediately placed in a sealed bag.

Water sampling was conducted at a water site in the Forest of Toyota on the same days as those of the ground surface sampling. Water was collected for 250 mL in total using a 20 mL syringe (Terumo, Japan) and filtered using a syringe filter made of hydrophilic acetylcellulose (25 mm diameter; 5 µm pore size, Membrane Solutions, USA).

Both non-woven sheets and the filters of water samples were transported from the field to the laboratory under refrigerated conditions and stored at -80 °C. To prevent cross-contamination during the operation, the operator worn clean new nitrile gloves each time when separating the sheets of each sample and when handling different samples.

A total of 90 ground surface and 29 water samples were collected. The weather conditions during sampling were obtained from the Japan Meteorological Agency (https://www.jma.go.jp/jma/index.html Accessed March 06, 2023), and accordingly, sampling was conducted during daylight hours, when there was no rainfall (Supplementary Table S2). Of the 30 sampling events, sampling on November 11, 2021, we obtained ground surface samples only, while no water samples were collected.

### DNA extraction

After thawing the collected non-woven sheets at room temperature, 100 mL of nuclease-free water was added and the bag was shaken to suspend terrestrial samples in water. Subsequently, the suspended water was filtered using a fresh 20 mL syringe and syringe filter made of hydrophilic acetylcellulose (25 mm diameter, 5 µm pore size. Membrane Solutions, USA) until all the sample volume was filtered or the filter was clogged. A maximum of two filters per sample was used. The filtration volume for each sample ranged from 35 mL to 80 mL. DNA was extracted using a NucleoSpin Soil Kit (Macherey-Nagel, Germany) according to the manufacturer’s instructions. Finally, 100 µL DNA was eluted for each sample.

### Library preparation and amplicon sequencing

The first polymerase chain reaction (PCR) was performed in 12 µL reaction volume containing 6.0 µL 2×KAPA HiFi HotStart ReadyMix (KAPA Biosystems, USA), 1.4 µL MiBird^28^ or MiMammal^12^ primers (5 µM primer F/R each), 2.6 µL nuclease-free water, and 2.0 µL environmental DNA sample. The temperature cycling after the first 3 min of denaturation at 95 °C was as follows: denaturation at 98 °C for 20 s, annealing at 65 °C for 15 s, and extension at 72 °C for 15 s for 35 cycles, followed by an additional extension at 72 °C for 5 min. The first PCR was performed in four replicates with the same sample to reduce PCR bias, and the amplicons were pooled after PCR. The pooled PCR products were purified using 0.8× AMPure XP (Beckman Coulter, USA) according to the manufacturer’s instructions. The purified PCR products were quantified using the Quantus Fluorometer and QuantiFluor ONE dsDNA System (Promega, Japan) to confirm amplification. The second PCR was performed in 25 µL reaction volume containing 12.5 µL 2× KAPA HiFi HotStart ReadyMix, 2.5 µL each primer of Nextera XT Index Kit v2 (Illumina, USA), 5 µL nuclease-free water, and 2.5 µL purified 1st PCR product. Different index combinations were used for each template. The temperature cycling after the first 3 min of denaturation at 95 °C was as follows: denaturation at 98 °C for 20 s and extension at 72 °C for 15 s for 10 cycles, followed by the final extension at 72 °C for 5 min. The second PCR product was purified using 0.8× AMPure XP according to the manufacturer’s instructions. The purified PCR products were quantified using the Quantus Fluorometer and QuantiFluor ONE dsDNA System (Promega, Japan). Subsequently, diluted to the same concentration, mixed in equal volumes, and measured using the Quantus Fluorometer and the QuantiFluor ONE dsDNA System. The library size was confirmed using the TapeStation and the D1000 kit (Agilent, USA), and the product was diluted to 1 nM concentration. Later, the product was diluted to a final concentration of 50 pM and sequenced for 150 bp paired-end using the iSeq 100 (Illumina, USA) with 20% PhiX spike-in.

### Sequence analysis

Illumina paired-end sequencing data were demultiplexed based on barcode sequences using the Local Run Manager (Illumina) in the iSeq 100 system and processed using QIIME2 (version 2019.1)^41^. The reads were cleaned of noise and primer sequences using the q2-dada2 plugin^42^. Forward and reverse sequences were merged to remove chimeric sequences, and a feature table and representative sequences were generated. Compared to the systems that use conventional flow cells, systems using patterned flow cells, including iSeq 100, have a higher frequency of index hopping, in which sequences from other samples contaminate the sequence information of the obtained samples, and^43^. To remove the read counts possibly caused by index hopping, a cutoff value for each representative sequence was set to 0.1% of the total read count of the ASV per run. The read count of the representative sequence of a sample was changed to zero when the count was smaller than the cutoff value. The generated representative sequences were searched using BLASTN against the NCBI database (https://ftp.ncbi.nlm.nih.gov/blast/db/v5/ Accessed November 05, 2021). Sequences classified as birds and mammals with 97% or higher identity were extracted. The results were then listed at the genus level.

To reduce detection loss in the database-matching phase, we conducted sequence confirmation for 17 representative bird and mammal species observed in the Forest of Toyota, which do not have sequences of MiBird and MiMammal regions registered in the National Center for Biotechnology Information (NCBI). To date, two species, the Japanese giant flying squirrel (*Petaurista leucogenys*) and pale thrush (*Turdus pallidus*), have not been recorded. For Japanese giant flying squirrels, the target sequence of the MiMammal region was confirmed from two samples of nest materials and two fecal samples collected near the nest box. The pale thrush was subjected to target sequence confirmation using axes from two different feather samples collected on different days in 2008 (Supplementary Table S3). The 170 bp sequence of the Japanese giant flying squirrel identified in this study showed 92.26% homology to that of *Petaurista hainana*, the closest NCBI-registered sequence, and the 182 bp sequence of the pale thrush showed 100% homology to those of *Turdus kessleri*, *Turdus obscurus*, *Turdus celaenops*, and *Turdus abyssinicus*. The representative sequences were searched against these two species and considered as detected if sequences with 97% or more homology with either of the sequences.

Although human DNA was often detected in the samples, likely because humans routinely visiting around the sampling area, representative sequences classified in the genus *Homo* were excluded from the downstream analysis, to limit our investigations on detecting wildlife.

Also, *Meleagris*, *Gallus*, *Coturnix*, *Bos*, *Capra*, and *Ovis* spp. are domesticated species that have never been observed in the Forest of Toyota, suggesting that their DNA had been contaminated from artificial resources. Furthermore, these species have been unexpectedly detected in the analysis of natural environments not only in Japan but also in other regions worldwide^16,26,27^. As we targeted wild animals, representative sequences classified as these six genera were excluded from the downstream analysis. Upset plots were created using UpSetR^44^ to visualize the number of bird and mammal species shared among the four sampled sites.

## Acknowledgements

The authors thank member of Social Inclusion Dept. Corporate Citizenship Div. Toyota Motor Corporation and the Forest of Toyota staff for their invaluable help in conducting monitoring at the Forest of Toyota and in the morphological identification of the animals.

## Author contributions

M.K. Conceptualization, Methodology, Fieldwork, Experiment, Data analysis, Writing (original draft), Writing (review & editing)

Y.F. Data analysis, Writing (review & editing)

H.T. Research Management, Fieldwork, Writing (review & editing) All authors discussed the results and commented on the manuscript.

## Data availability

Sequencing data generated during this study is available on the DDBJ Sequence Read Archive under accession numbers PRJDB18462.

## Supplementary Information

**Supplementary Table S1:**
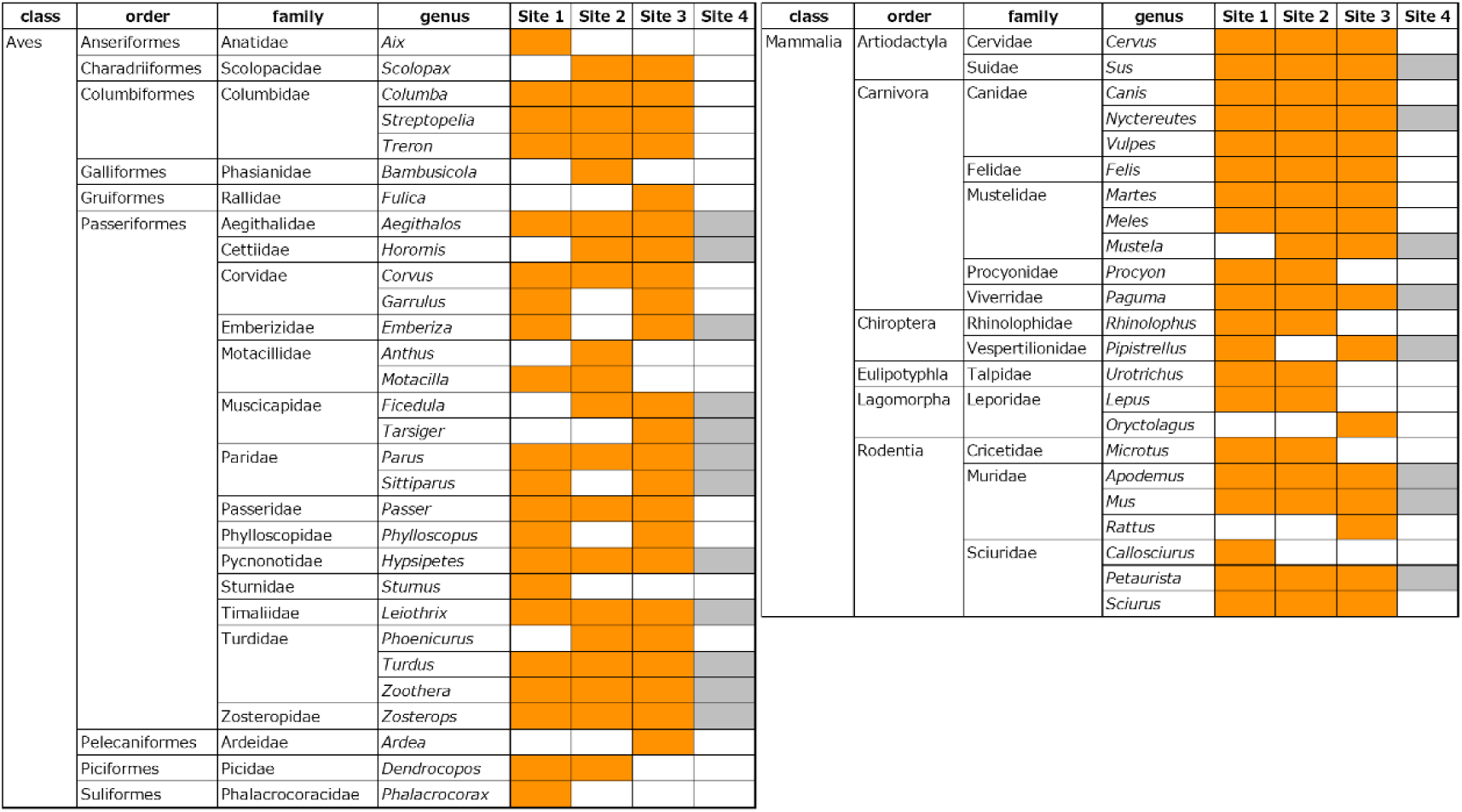
The location where bird and mammal DNA was detected.

**Supplementary Table S2:**
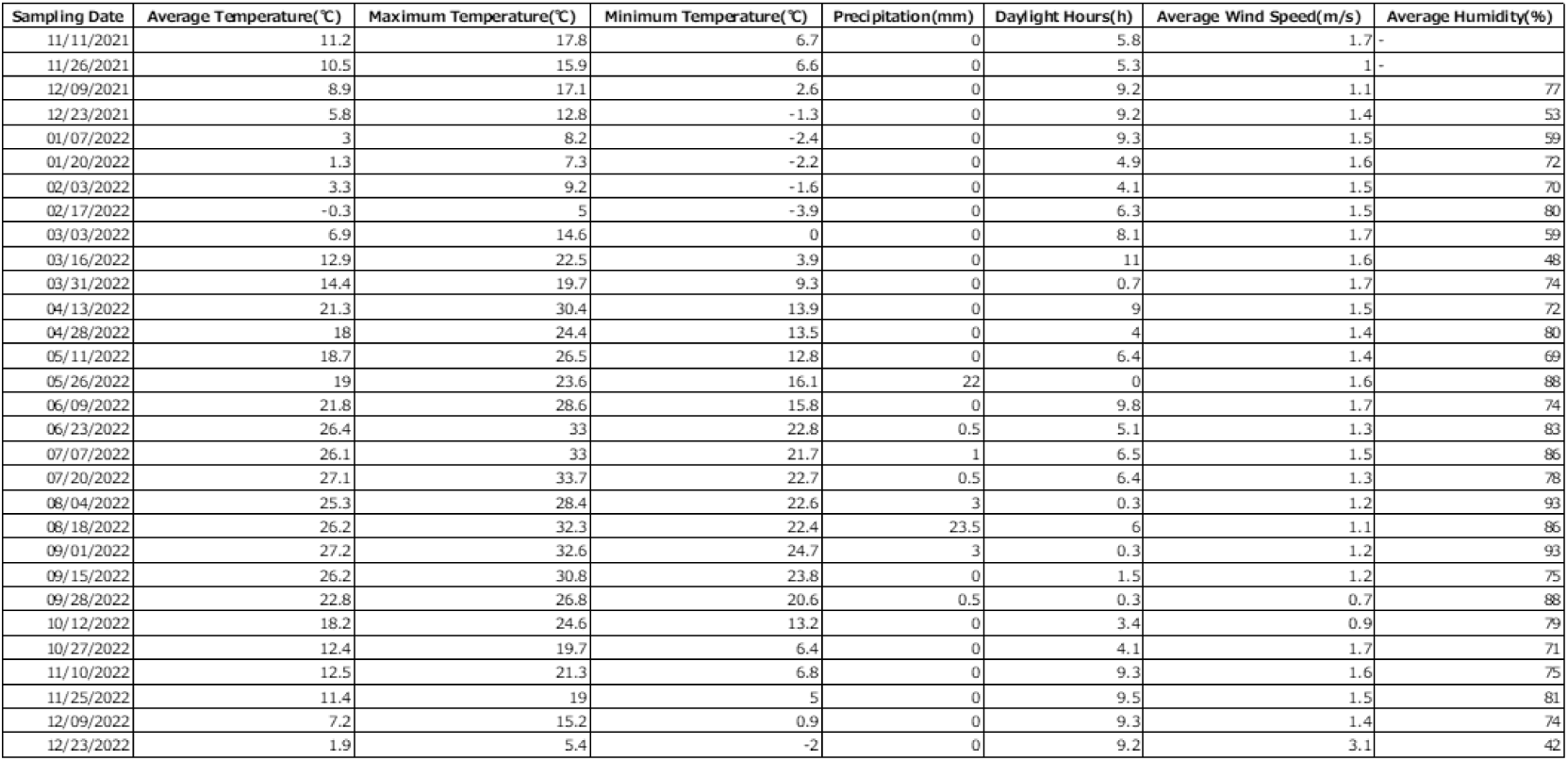
Details of when the DNA sampling was conducted and the weather conditions on that day.

**Supplementary Table S3:**
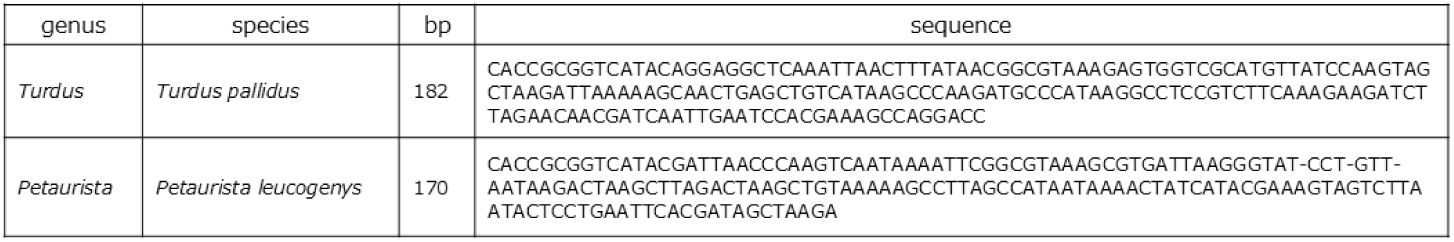
DNA sequences obtained from biological samples separately from this experiment.

